# *Pax9* is required for cardiovascular development and interacts with *Tbx1* in the pharyngeal endoderm to control 4^th^ pharyngeal arch artery morphogenesis

**DOI:** 10.1101/576660

**Authors:** Helen M. Phillips, Catherine A. Stothard, Wasay Mohiuddin Shaikh Qureshi, Anastasia I. Kousa, J. Alberto Briones-Leon, Ramada R. Khasawneh, Rachel Sanders, Silvia Mazzotta, Rebecca Dodds, Kerstin Seidel, Timothy Bates, Mitsushiro Nakatomi, Simon J. Cockell, Jürgen E. Schneider, Timothy J. Mohun, René Maehr, Ralf Kist, Heiko Peters, Simon D. Bamforth

## Abstract

Developmental defects affecting the heart and aortic arch arteries are a key phenotype observed in DiGeorge syndrome patients and are caused by a microdeletion on chromosome 22q11. Heterozygosity of *TBX1*, one of the deleted genes, is expressed throughout the pharyngeal arches and is considered a key component for the arch artery defects. *Pax9* is expressed in the pharyngeal endoderm and is downregulated in *Tbx1* mutant mice. We show here that *Pax9* deficient mice are born with complex cardiovascular malformations affecting the outflow tract and aortic arch arteries with failure of the 3^rd^ and 4^th^ pharyngeal arch arteries to form correctly. Transcriptome analysis indicated that *Pax9* and *Tbx1* may function together, and mice double heterozygous for *Tbx1/Pax9* presented with a significantly increased incidence of interrupted aortic arch when compared to *Tbx1* heterozygous mice. Using a novel *Pax9Cre* allele we demonstrated that the site of this *Tbx1-Pax9* genetic interaction is in the pharyngeal endoderm, therefore revealing that a *Tbx1/Pax9*-controlled signalling mechanism emanating from the pharyngeal endoderm is required for critical tissue interactions during normal morphogenesis of the pharyngeal arch artery system.

**Summary statement:** *Pax9* is required for outflow tract and aortic arch development, and functions together with *Tbx1* in the pharyngeal endoderm for 4^th^ arch artery formation.

## Introduction

Conotruncal heart malformations, affecting the outflow tract and aortic arch arteries, occur in 30% of all cases of congenital heart defects (Thom et al., 2006) and are a major cause of morbidity and death. Approximately 20% of foetuses identified with conotruncal defects by ultrasound are diagnosed with 22q11 deletion syndrome (22q11 DS) (Boudjemline et al., 2001), the most common microdeletion syndrome with an incidence of 1:4000 live births (Scambler, 2000). Patients typically have a 3Mb deletion on chromosome 22 that encompasses 45 protein coding genes (Morrow et al., 2018) and ~80% of patients present with some form of congenital cardiovascular defect (Momma, 2010), of which one of the most common observed is interruption of the aortic arch (Unolt et al., 2018). Although clinically rare in the general population, approximately 50% of all cases of interrupted aortic arch occur in 22q11 DS patients (Boudjemline et al., 2001; Lewin et al., 1997; Van Mierop and Kutsche, 1986). Interrupted aortic arch is a consequence of the left 4^th^ pharyngeal arch artery (PAA) failing to form correctly during embryonic development. There are five pairs of PAA that arise within a series of repeated protuberances on either side of the developing pharynx known as the pharyngeal arches. These arches consist of paraxial mesoderm- and neural crest cell (NCC)-derived mesenchyme, and are externally enclosed by ectoderm and internally lined by endoderm (Graham, 2003). The initially bilaterally symmetrical PAA develop sequentially in a cranial to caudal sequence during embryogenesis but undergo a complex remodelling process that is conserved in mammals to form the asymmetrical mature aortic arch configuration (Bamforth et al., 2013; Hiruma et al., 2002). Of the genes deleted in 22q11 DS, hemizygosity of *TBX1* is considered to be the cause of the cardiovascular defects seen in these patients (Jerome and Papaioannou, 2001; Lindsay et al., 2001; Merscher et al., 2001). Tbx1 is expressed throughout the pharyngeal arches within the ectoderm, endoderm and mesoderm cells and is critical for pharyngeal arch and PAA development. Complete loss of *Tbx1* from the mouse results in a failure of the pharyngeal arches to form correctly resulting in hypoplasia of the second arch and aplasia of the remaining caudal arches, leading to an absent thymus and common arterial trunk (Jerome and Papaioannou, 2001; Merscher et al., 2001). In contrast, heterozygous deletion of *Tbx1* only affects development of the 4^th^ PAA resulting in interruption of the aortic arch in a minority of mice with aberrant right-subclavian artery observed more frequently (Lindsay et al., 2001). Although almost all 22q11 DS patients are hemizygous for the deletion, the phenotypic spectrum is highly variable (Unolt et al., 2018), thus it has been proposed that genes outside of the deleted region may impact on the clinical phenotype (Guo et al., 2011). Several genes have been identified as potential modifiers of 22q11 DS as determined by genetic interaction studies and transcriptome analyses in *Tbx1* mutant mice (Aggarwal and Morrow, 2008; Ivins et al., 2005; Liao et al., 2008; Papangeli and Scambler, 2013). One such candidate gene is *Pax9* whose expression in the pharyngeal endoderm was found to be significantly reduced in Tbx1-deficient embryos (Ivins et al., 2005; Liao et al., 2008), although further investigations into the nature of any putative interaction have not been reported. *Pax9* belongs to a family of Pax genes encoding for transcription factors involved in various cellular roles during embryonic development (Chi and Epstein, 2002; Mansouri et al., 1996) and is specifically expressed in the endoderm of all four pharyngeal pouches of the mouse by E9.5 (Neubuser et al., 1995). At later stages in mouse development, Pax9 is expressed in the oesophagus, somites and limbs, and within the neural crest contributing to craniofacial development (Peters et al., 1998). Embryos deficient for *Pax9* exhibit craniofacial defects, including cleft secondary palate as well as absent teeth, skeletal defects, and lack derivatives of the 3^rd^ and 4^th^ pharyngeal pouches, i.e. the thymus, parathyroids and ultimobranchial bodies (Peters et al., 1998) yet the role of Pax9 in cardiovascular development has not been determined.

In this study we have investigated cardiovascular development in Pax9-deficient mice, and show that loss of *Pax9* leads to complex cardiovascular defects affecting the outflow tract and aortic arch arteries. We also uncover a strong genetic interaction between *Tbx1* and *Pax9* that leads to 4^th^ PAA-derived defects in double heterozygous mice, and this interaction is cell-autonomous within the pharyngeal endoderm.

## Results

### *Pax9*-deficient embryos have cardiovascular defects

Mice lacking *Pax9* die perinatally and exhibit craniofacial, pharyngeal gland and skeletal defects (Peters et al., 1998). A cleft secondary palate had initially been suggested as a cause for postnatal lethality of *Pax9*-deficient mice, however, recent observations showed that ~1 *%* of *Pax9−/−* mice have an intact secondary palate but still die shortly after birth (data not shown). We therefore carefully re-analysed *Pax9^−/−^* neonates from a *Pax9^+/−^* intercross to identify the cause of death. When dissected, we found that all *Pax9^−/−^* neonates had a patterning defect of their aortic arch arteries which included an interrupted aortic arch with aberrant right subclavian artery (**Fig. 1B; Table 1**). To investigate these defects in more detail we processed *Pax9^−/−^* embryos for histology and magnetic resonance imaging (MRI) and found that *Pax9^−/−^* embryos presented with a wide range of cardiovascular abnormalities (**Fig. 1C-Q; Table 1**). These defects included perimembranous ventricular septal defect (VSD), double-outlet right ventricle with interventricular communication, interruption of the aortic arch and aberrant right subclavian artery. The common carotid arteries were also frequently absent resulting in unilateral or bilateral internal and external carotids arising directly from the dorsal aorta and the aberrant right subclavian artery. *Pax9^−/−^* embryos also had a hypoplastic aorta (**Fig. 1M-O**) and bicuspid aortic valves (**Fig. 1Q**).

**Figure 1.**
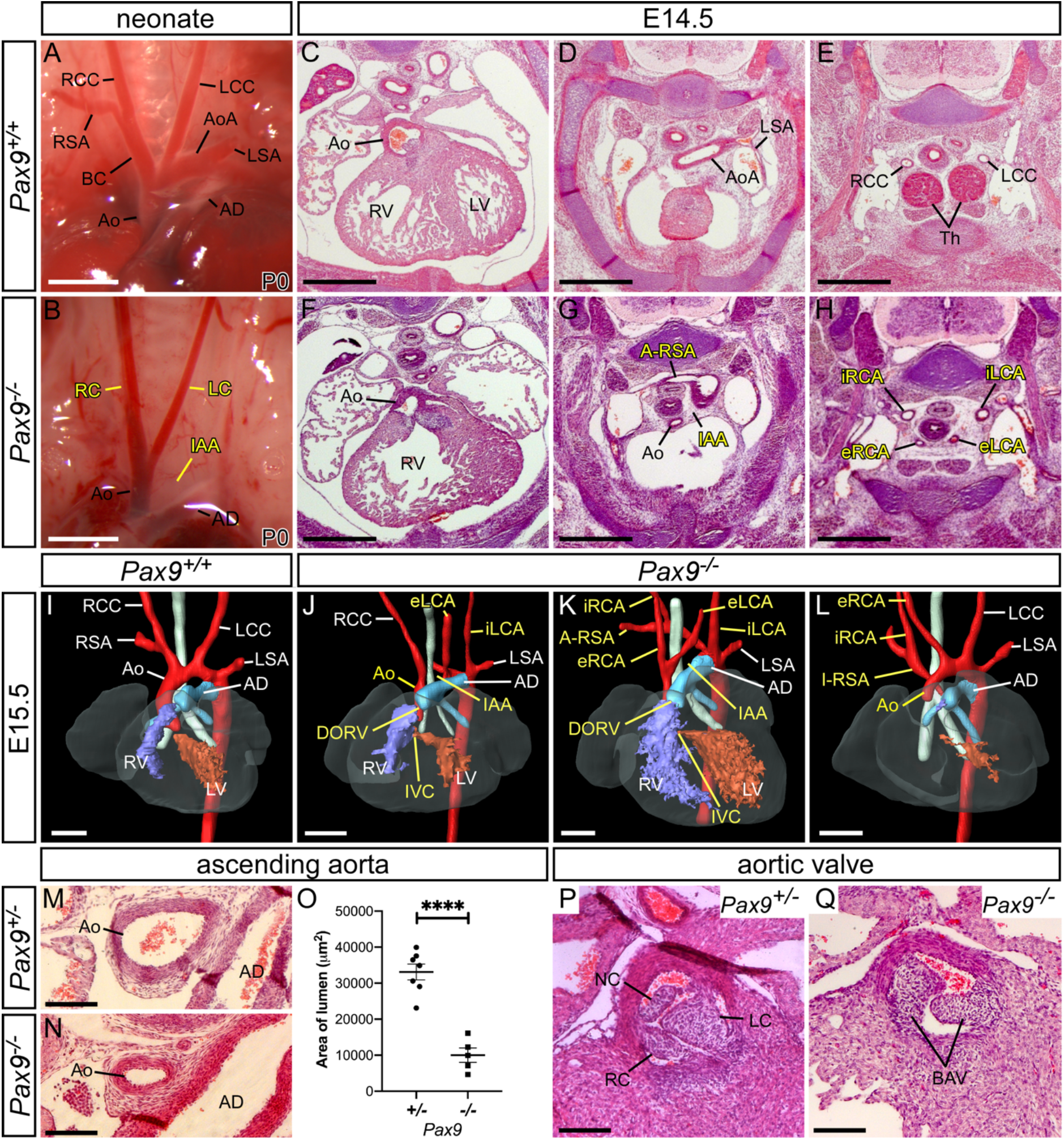
Loss of *Pax9* results in cardiovascular developmental defects. **A**, Arch arteries of control neonates were normal (n=8). **B**, *Pax9^−/−^* neonates (n=5) displayed interrupted aortic arch (IAA), absent right subclavian artery (presumed retro-esophageal), and atypical right and left carotid arteries (LC, RC). Scale, 1mm. **C-H**, E14.5 histology. **C-E**, *Pax9^+/+^* embryos show normal outflow tract, arch arteries and thymus. **F-H**, In *Pax9^−/−^* embryos (n=4) the aorta (Ao) arises aberrantly from the right ventricle giving double-outlet right ventricle (DORV; **F**). The aorta is hypoplastic and interrupted (IAA), and an aberrant right subclavian artery (A-RSA) is present (**G**). More cranially the thymus is absent and aberrant internal and external carotid arteries (iRCA, eRCA, iLCA, eLCA) are seen. **I-L**, 3-D reconstructions of E15.5 hearts from MRI datasets. **I**, *Pax9^+/+^* embryo with normal aortic arch arteries. **J-L**, In Pax9^−/−^ embryos (n=15) defects seen are DORV with interventricular communication (IVC), IAA, A-RSA (retro-esophageal in **J, K** and isolated in **L**) and aberrant carotid arteries. Scale, 500μm. **M-O**, The ascending aorta is significantly smaller in *Pax9^−/−^* embryos (n=5) compared to *Pax9^+/−^* control embryos (n=7). Mean ± s.e.m.; ****p<0.0001; two-tailed unpaired t-test. **P**, *Pax9^+/−^* control embryos have three normal aortic valve leaflets (n=6), the right (RC), left (LC) and non-coronary (NC). **Q**, *Pax9^−/−^* embryos (n=5) have bicuspid aortic valve. Scale, 100μm. Abbreviations: Ao, aorta; AoA, aortic arch; AD, arterial duct; BC, brachiocephalic; dAo, dorsal aorta; LCC, left common carotid artery; LSA, left subclavian artery; LV, left ventricle; RCC, right common carotid artery; RSA, right subclavian artery; RV, right ventricle; Th, thymus.

**Table 1.**
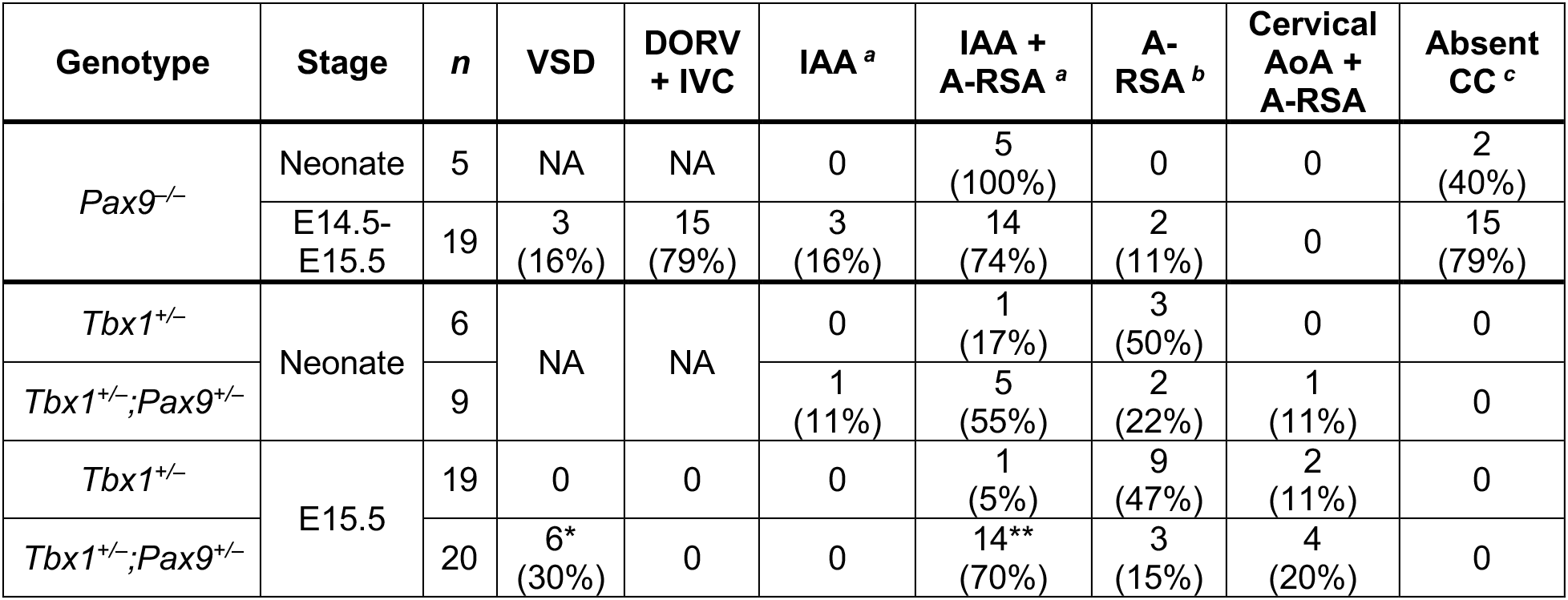
Cardiovascular defects observed in mutant embryos (>E14.5) and neonates. *Pax9^−/−^* neonates (n=5) were analysed by dissection, and embryos at E14.5 by histology (n=4) and at E15.5 by MRI (n=15). *Tbx1^+/−^* (n=19) and *Tbx1^+/−^;Pax9^+/r^* (n=20) embryos were analysed by MRI. All wild-type (n=16) and *Pax9^+/−^* (n=15) embryos assessed were normal. There was a statistically significant increase in the incidence of VSD and IAA in the *Tbx1^+/−^;Pax9^+/−^* embryos compared to the *Tbx1^+/−^* embryos; *p<0.05; **p<0.0001; Pearson’s chi-squared test for associations.^*a*^ Combined incidence of interrupted aortic arch (IAA), irrespective of co-occurrence with aberrant right subclavian artery (A-RSA), in *Pax9^−/−^* neonates and embryos (n=24), was 92% (n=19 type-B and n=3 type-C). ^*b*^ A-RSA refers to a retroesophageal, cervical origin or isolated right subclavian artery. Two *Pax9^−/−^* embryos had isolated RSA. ^c^ Absent common carotid artery (CC), resulting in the internal and external carotid arteries arising directly from the main aortic vessels, unilaterally (n=4) or bilaterally (n=11) in *Pax9^−/−^* embryos. Abbreviations: AoA, aortic arch; DORV + IVC, double-outlet right ventricle with interventricular communication; VSD, perimembranous ventricular septal defect; NA, not assessed.

Developmental defects of the arch arteries occur when the PAA fail to form or remodel correctly. To investigate the morphogenesis of the PAA we performed high-resolution episcopic microscopy (HREM) on E10.5 and E11.5 *Pax9^−/−^* embryos and manually segmented the datasets to create 3-D images of the PAA. Analysis of E10.5 coronal sections revealed that the 3^rd^ and 4^th^ pharyngeal arches of *Pax9^−/−^* embryos were smaller compared to *Pax9^+/+^* controls (**Fig. 2A-C**), as previously reported (Peters et al., 1998). At E10.5 the developing PAA are bilaterally symmetrical, having formed in a cranial to caudal sequence. The 1^st^ and 2^nd^ PAA have normally regressed by this stage, and the 3^rd^, 4^th^ and 6^th^ PAA are present (Bamforth et al., 2013; Hiruma et al., 2002) (**Fig. 2A**). In *Pax9^−/−^* embryos the 1^st^ and 2^nd^ PAA were abnormally persistent, the 3^rd^ PAA was either absent, hypoplastic or interrupted, and the 4^th^ PAA was absent (**Fig. 2B, C; Table 2**). The aortic sac was also hypoplastic in *Pax9^−/−^* embryos compared to the controls (**Fig. 2B, C**). Intra-cardiac injection of India ink at E10.5-E11.0 showed that, in controls, the caudal PAA were patent to ink and of equivalent size (**Fig. 2D**). In *Pax9^−/−^* embryos, however, persisting 1^st^ and 2^nd^ PAA were patent to ink, and the 3^rd^ PAA were hypoplastic (**Fig. 2J – L**) or non-patent to ink, and therefore presumed absent. The majority of the 4^th^ PAA were bilaterally non-patent to ink (**Fig. 2J – L; Table 2**). Immunohistochemical staining for Pecam1 showed that the endothelium within the 3^rd^ and 4^th^ pharyngeal arches had formed lumenised PAA at E10.5 in control embryos (**Fig. 2G**), but in *Pax9^−/−^* embryos the 3^rd^ PAA was visibly smaller in size and only isolated endothelial cells were seen within the 4^th^ pharyngeal arch (**Fig. 2H, I**).

**Figure 2.**
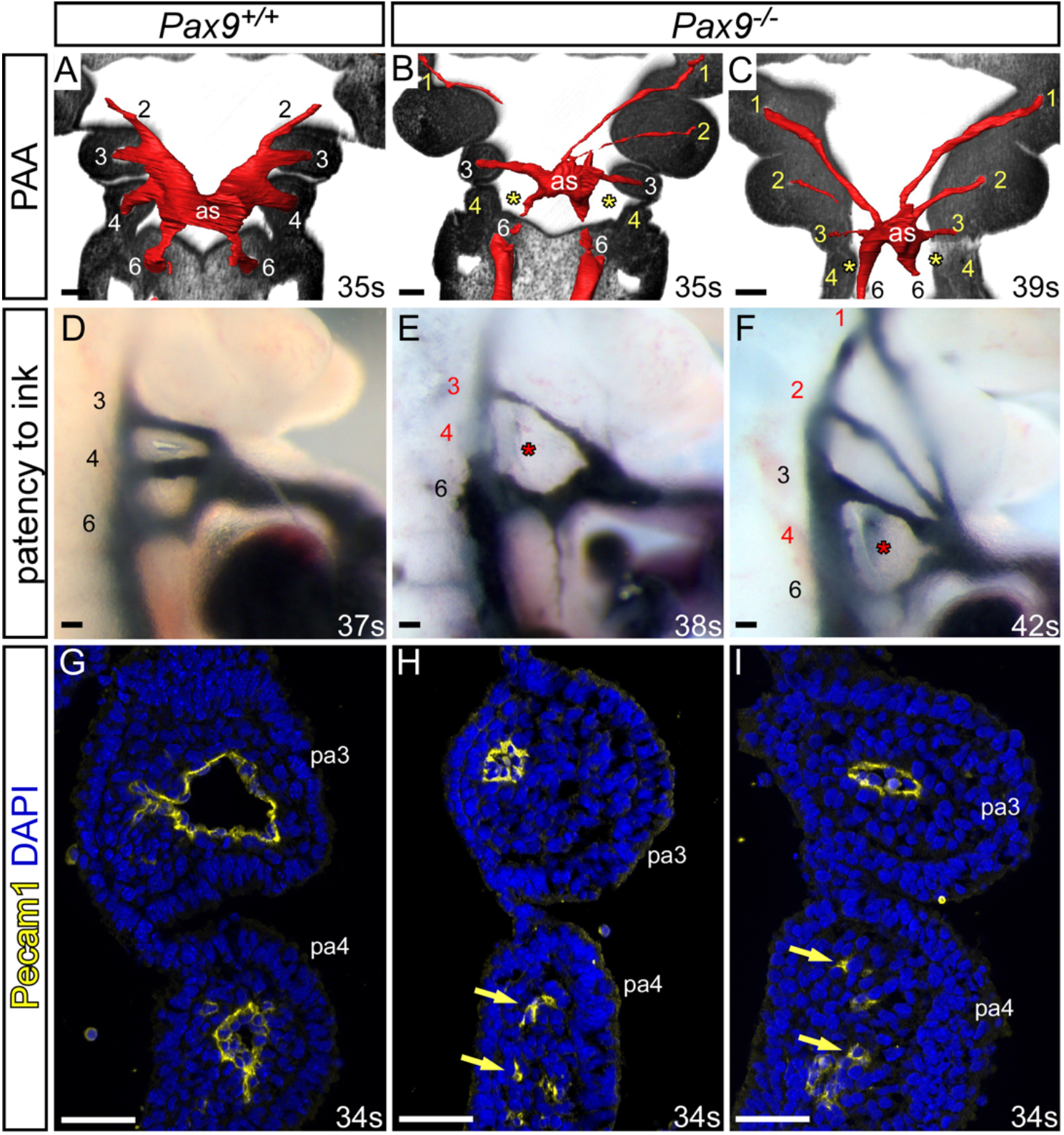
*Pax9* loss causes aberrations in pharyngeal arch artery formation. **A-C**, Coronal views of control *(Pax9^+/+^)* and mutant *(Pax9^−/−^)* embryos at E10.5 (30-40s), examined by high-resolution episcopic microscopy. **A**, In *Pax9^+/+^* control embryos (n=5) the 3^rd^, 4^th^ and 6^th^ PAA are of equal size and bilaterally symmetrical. **B, C**, In *Pax9^−/−^* embryos (n=4) the 4^th^ PAA was bilaterally absent *(asterisk)*, the 3^rd^ PAA and aortic sac (as) were hypoplastic, and the 1^st^ and/or 2^nd^ PAA abnormally persisted. **D-F**, Intracardiac ink injection into E10.5-E11.0 (31-42s) embryos. **D**, In control embryos (n=12) PAA 3-6 are patent to ink, are of equivalent diameter and bilaterally symmetrical. **E, F**, In *Pax9^−/−^* embryos (n=12) the 3^rd^ PAA is hypoplastic and the 4^th^ PAA non-patent to ink *(asterisk).* **F**, The 1^st^ and 2^nd^ PAA are persisting anomalously. **G-I**, Immunostaining using Pecam1 antibody at E10.5 (31-36s). **G**, Control embryos (n=3) have a ring of Pecam1 labelled endothelium lining the 3^rd^ and 4^th^ PAA. **H, I**, In *Pax9^−/−^* embryos (n=5) the 3^rd^ PAA is visibly smaller and disorganized endothelial cells are within the 4^th^ pharyngeal arch (pa; *arrows).* Somite counts (s) indicated. Scale, 100μm.

**Table 2.**
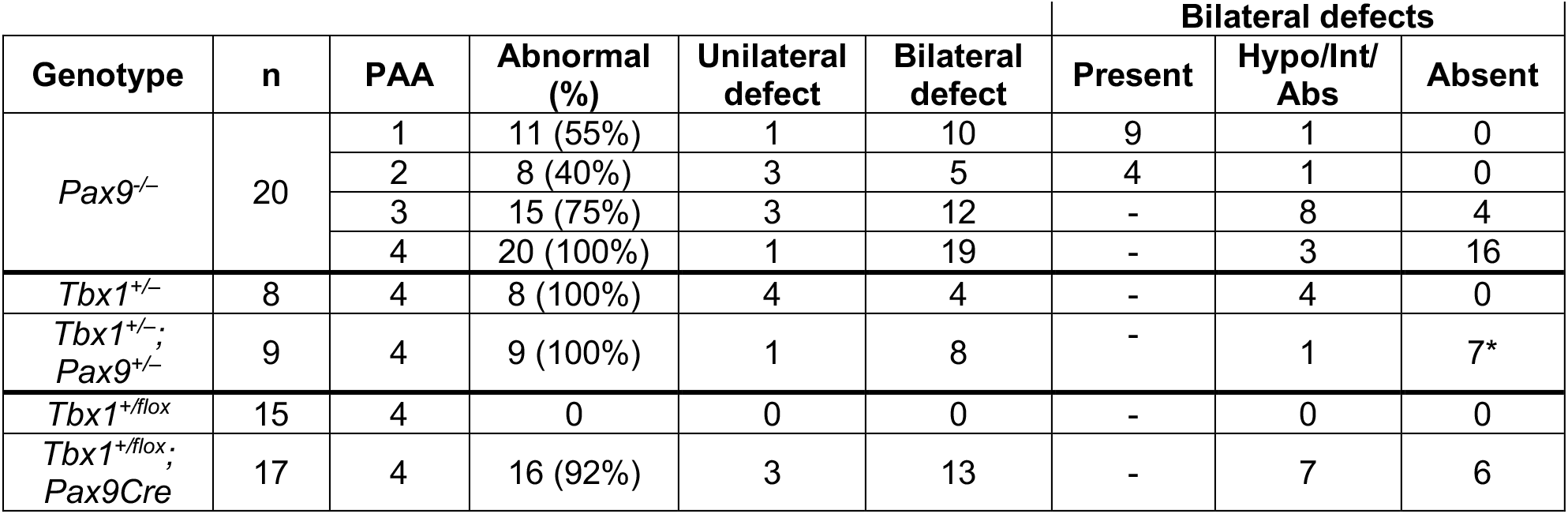
Pharyngeal arch artery defects. Embryos were collected and assessed for PAA defects. *Pax9^−/−^* embryos were either injected intra-cardially with ink (E10.5; n=12) or imaged using HREM (E10.5-E11.5; n=8) to visualise the PAA. Data are combined. For *Pax9^−/−^* embryos each left and right PAA 1-4 was scored as having a unilateral or bilateral defect, and the bilateral defects categorised as either present, a combination of hypoplastic, interrupted and/or absent (Hypo/Int/Abs), and bilaterally absent. For *Tbx1;Pax9* genotypes ink injection was performed at E10.5 and only the 4^th^ PAA was scored. The increase in bilaterally absent 4^th^ PAA in *Tbx1^+/−^;Pax9^+/−^* embryos is significant compared to *Tbx1^+/−^* embryos; *p=0.004; Pearson’s chi-squared test for associations. All control embryos, *Pax9^+/+^* (n=18), *Pax9^+/−^* (n=25), were normal.

During E11.5 of mouse development, the PAA system begins to remodel and becomes asymmetric in appearance (Bamforth et al., 2013; Hiruma et al., 2002). In *Pax9* control embryos at this stage the right 6^th^ PAA appeared thinner and the distal outflow tract was septated. The carotid duct, the region of the dorsal aorta between the 3^rd^ and 4^th^ PAA, had begun to involute but the 3^rd^, 4^th^ and 6^th^ PAA remained connected to the dorsal aortae (**Fig. 3A**). In *Pax9^−/−^* embryos at E11.5, evidence of presumed persisting 1^st^ or 2^nd^ PAA were observed extending anteriorly from the aortic sac, and the 3^rd^ PAA were also affected, either unilaterally or bilaterally, and were either absent, hypoplastic or interrupted and not connected to the dorsal aortae (**Fig. 3B-D**). The carotid ducts were maintained, and this, coupled with the abnormal detachment of the 3^rd^ PAA from the dorsal aorta and the persistence of the 1^st^ or 2^nd^ PAA, resulted in an absent common carotid artery and the internal and external carotid arteries arising directly from the aortic arch arteries as observed at E15.5 (**Fig. 1**). Additionally, the distal outflow tract was unseptated and the 4^th^ PAA were bilaterally absent (**Fig. 3B-D; Table 2**).

**Figure 3.**
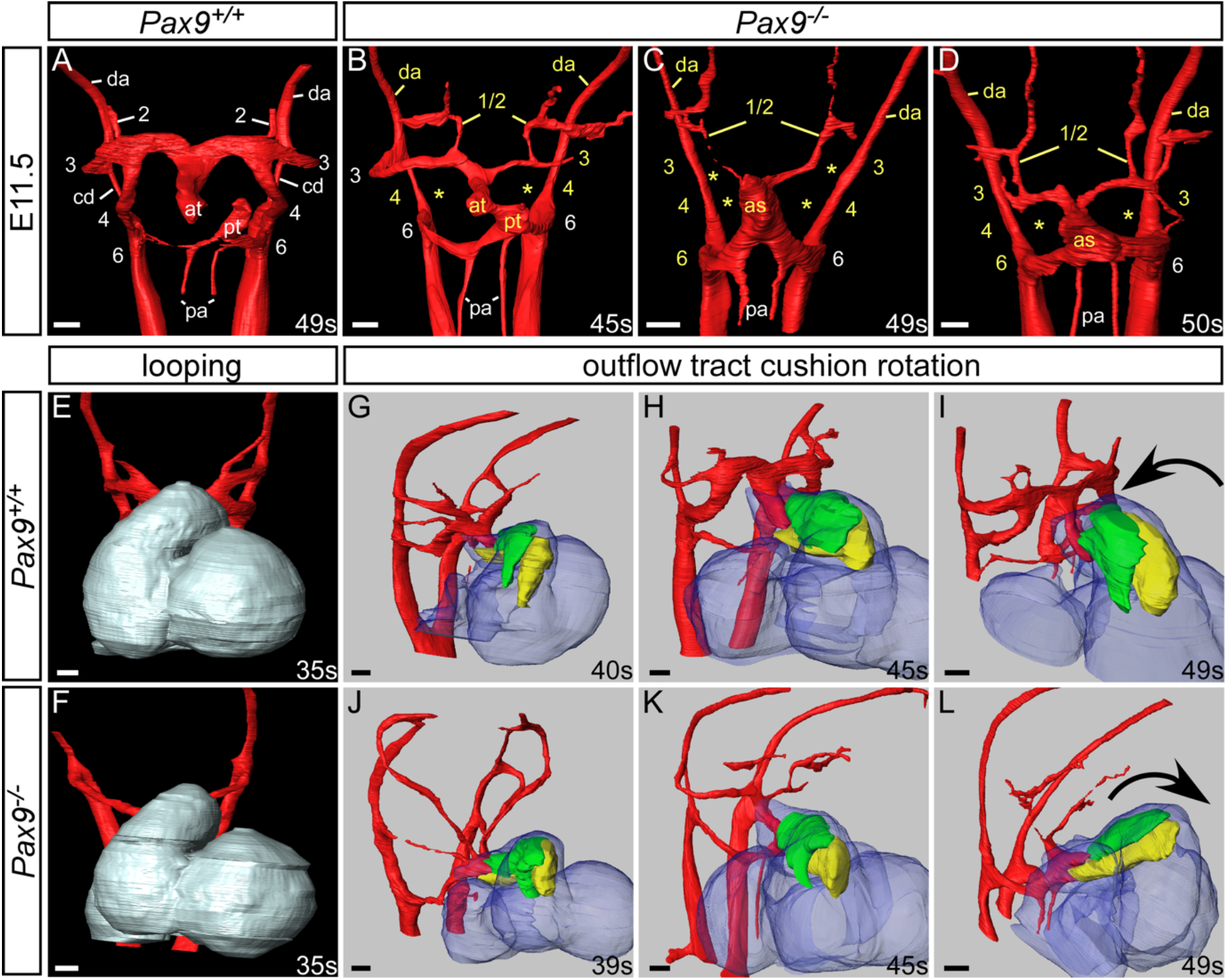
*Pax9^−/−^* embryos display aberrant rotation of the outflow tract and abnormal PAA remodelling. Image datasets were acquired by high-resolution episcopic microscopy. **A-D**, 3-D reconstructions of the PAA at E11.5 (45-50s). **A**, In *Pax9^+/+^* control embryos (n=5) the outflow tract is septated, the right 6^th^ PAA has thinned and the carotid duct (cd) has begun to involute. **B-D**, In *Pax9^−/−^* embryos (n=4) the 4^th^ PAA are absent and the 3^rd^ PAA are aberrantly connected or detached from the dorsal aorta *(asterisks).* The carotid duct has failed to involute and septation of the outflow tract is delayed. Persisting 1^st^ or 2^nd^ PAA are observed coursing anteriorly from the abnormal 3^rd^ PAA (**B, D**) or from the aortic sac **(C). E, F**, Looping of the heart tube at E10.5 was not affected in *Pax9^−/−^* embryos (n=4). G-L, Outflow tract rotation from E10.5 to E11.5 (39-49s). **G-I**, In control embryos the major outflow tract cushions (parietal in *green;* septal in *yellow*) rotated in an anti-clockwise direction resulting in the cushions being aligned side-by-side. **J-L**, In *Pax9^−/−^* embryos the cushions rotated in a clockwise direction resulting in the parietal cushion anterior to the septal cushion. Abbreviations: as, aortic sac; at, aortic trunk; da, dorsal aorta; pa, pulmonary arteries; pt, pulmonary trunk. Somite (s) counts indicated. Scale, 100μm.

Looping of the heart tube at E10.5 was not affected in *Pax9^−/−^* embryos (**Fig. 3E, F**). In normal mouse development the outflow tract cushions rotate through an anticlockwise direction (Bajolle et al., 2006) resulting in the major septal and parietal cushions being positioned side by side (**Fig. 3G-I**). In *Pax9^−/−^* embryos, however, by the end of the E11.5 stage these cushions were aberrantly positioned, with the parietal cushion more anterior to the septal cushion, indicating that a clockwise rotation of the outflow tract cushions had occurred (**Fig. J-L**). This failure in proper rotation of the major cushions is potentially the cause of the double-outlet right ventricle phenotype observed at E15.5.

This data demonstrates that *Pax9^−/−^* embryos display a range of cardiovascular developmental defects resulting from the abnormal morphogenesis of the PAA. The 1^st^ and 2^nd^ PAA often persist, the 3^rd^ PAA forms but is smaller and subsequently collapses, and the 4^th^ PAA fails to form at all. These defects directly lead to the absent common carotid artery, aberrant right subclavian artery and interrupted aortic arch phenotypes seen in foetal and neonatal stages in *Pax9^−/−^* mice. Moreover, delayed outflow tract septation occurs with abnormal rotation leading to the double-outlet right ventricle phenotype.

### Pharyngeal endoderm signalling influences neural crest cell differentiation

During normal embryogenesis, a subset of NCC, the cardiac neural crest, migrate into the caudal pharyngeal arches to provide support to the developing PAA and also contribute to outflow tract septation (Hutson and Kirby, 2003). *Pax9* is predominantly expressed in the pharyngeal endoderm (**Fig. S1**), but is also known to be expressed within the NCC contributing to craniofacial development (Kist et al., 2007), although expression in cardiac NCC has not been described. We therefore looked to see if a cell autonomous loss of *Pax9* from NCC resulted in cardiovascular defects. Conditional deletion of *Pax9* using *Wnt1Cre* transgenic mice (Danielian et al., 1998), and the *Pax9^flox^* allele (Kist et al., 2007), however, did not result in cardiovascular abnormalities, although cleft palate was observed as expected (**Fig. 4A-D**). We next looked to see if the migration of NCC was affected in global *Pax9^−/−^* embryos using *Wnt1Cre* and *eYFP* (Srinivas et al., 2001) reporter mice to create embryos where the NCC could be lineage traced. Live fluorescence imaging of *Pax9^+/+^;Wnt1Cre;eYFP* embryos at E10.5 revealed NCC present in the head and dorsal route ganglia, as well as migrating towards and filling the pharyngeal arches (**Fig. 4E, G, I**). A similar pattern was observed in *Pax9^−/−^;Wnt1Cre;eYFP* embryos (**Fig. 4F, H, J**). Immunostaining for BrdU incorporation specifically in NCC showed no significant difference in NCC proliferation in the 4^th^ pharyngeal arch at E10.5 (**Fig. 4K-M**), and very little apoptosis occurred throughout the caudal pharyngeal arches in both control and *Pax9^−/−^* embryos (**Fig. 4N-P**). These observations indicated that the migration of NCC *per se* to the pharyngeal arches were not affected in *Pax9^−/−^* embryos.

**Figure 4.**
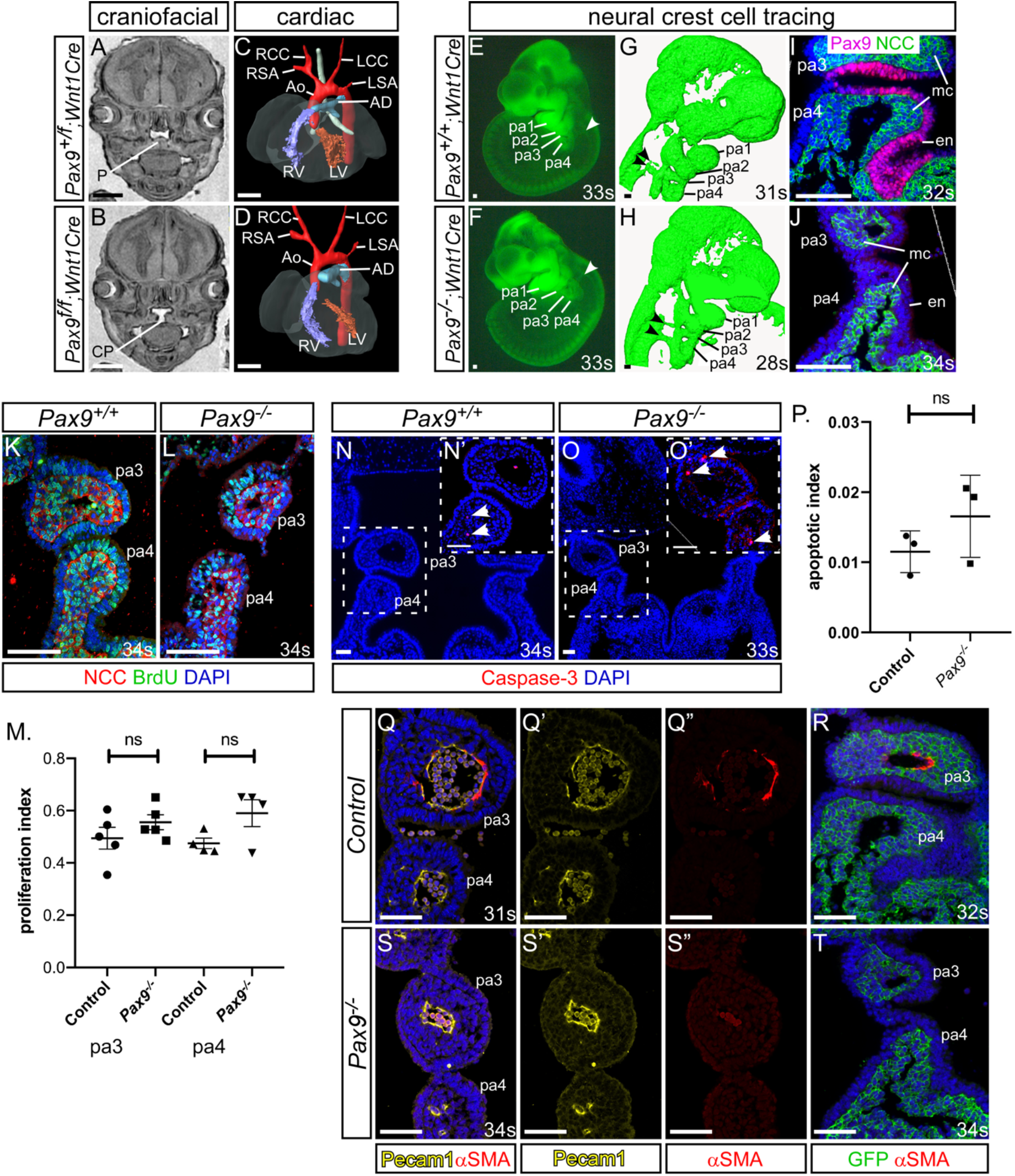
Failure in smooth muscle cell recruitment causes the 3^rd^ PAA to collapse. **A-D**, E15.5 embryos with a *Pax9* conditional deletion in neural crest cells (NCC) were examined by MRI. **A**, Control embryos had a normal palate (P). **B**, *Pax9^f/f^;Wnt1Cre* mutant embryos (n=6) had cleft palate (CP). **C, D**, No cardiovascular defects were observed in control or mutant embryos. Abbreviations as in Figure 1. **E-J**, NCC in *Pax9^+/+^* (**E, G, I**) or *Pax9^−/−^* (**F, H, J**) embryos were labelled using *Wnt1Cre* and *eYFP* alleles. **E, F**, Fluorescence imaging showed no difference in NCC migration into the pharyngeal arches *(arrowheads)* in control (n=7; 31-34s) and mutant embryos (n=5; 28-35s). **G, H**, Labelled NCC viewed by confocal microscopy also showed similar NCC migration patterns in control and mutant embryos *(arrowheads)*. **I, J**, Fluorescent embryos were sectioned coronally and immunostained using anti-GFP and anti-Pax9 antibodies. **J**, The caudal pharyngeal arches were smaller in *Pax9^−/−^* embryos. **K-L**, Embryo sections at E10.5 were immunostained with anti-BrdU and anti-GFP antibodies to detect proliferation in NCC in *Pax9^+/+^* (**K**) and *Pax9*^−/−^ (**L**) embryos at E10.5 (n=5 per genotype; 31-39s). **M**, No significant difference in the rate of proliferation was found between control and *Pax9^/–^* NCC. N-O, Embryo sections at E10.5 were immunostained with an anti-Caspase3 antibody to detect apoptosis in the caudal pharyngeal arches of *Pax9^+/+^* (**N**) and *Pax9^−/−^* (**O**) embryos at E10.5 (n=3 per genotype; 31-35s). Higher magnification views shown in dashed boxes (**N’, O’**). **P**, No significant difference in the rate of apoptosis was found between control and *Pax9^−/−^* embryos. Twotailed unpaired t-test. **Q-T**, E10.5 embryo sections were immunostained using anti-αSMA and anti-Pecam1 antibodies for smooth muscle and endothelium respectively (**Q, S**) or anti-αSMA and anti-GFP to label NCC in *Wnt1Cre;eYFP* embryos (**R, T**). In all control embryos smooth muscle cells surrounding the 3^rd^ PAA was observed (n=4; 31-36s; **Q, R**). In *Pax9^−/−^* embryos the 3^rd^ PAA had no visible recruitment of smooth muscle cells (n=6; 32-35s; **S, T**). Abbreviations: en, endoderm; mc, mesenchyme; pa, pharyngeal arch. Somite counts (s) indicated. Scales: A – D, 500μm; E-O, 100μm; Q-T, 50μm.

It therefore appears that the rapid collapse of the 3^rd^ PAA at E11.5 in *Pax9^−/−^* embryos, resulting in an absent common carotid artery, cannot be explained by aberrant NCC death, proliferation or migration pattern at E10.5. Given that NCC differentiate into the smooth muscle cells that support the remodelling PAA (Waldo et al., 1996), we looked to see if smooth muscle cell recruitment to the 3^rd^ PAA was affected in *Pax9^−/−^* embryos. Sections from E10.5 embryos were immunostained using an anti-alpha smooth muscle actin antibody to visualise smooth muscle cells around the 3^rd^ PAA. As expected, no staining was observed in control embryos around the 4^th^ PAA at this stage (Losa et al., 2017). Evidence of smooth muscle cell recruitment to the 3^rd^ PAA was seen in each control embryo examined (**Fig. 4Q, R**), however no evidence of any smooth muscle cell investment around the 3^rd^ PAA in *Pax9^−/−^* embryo sections was observed (**Fig. 4S, T**). These results indicated that the 3^rd^ PAA in *Pax9^−/−^* embryos forms but collapses due to a failure of smooth muscle cell investment, possibly from a perturbation in signals that emanate from the pharyngeal endoderm under Pax9 control.

### *Tbx1* and *Pax9* interact in 4^th^ PAA formation

The failure of the 4^th^ PAA to form in *Pax9^−/−^* embryos, however, is likely to be due to an alternative mechanism as NCC are not required for initial PAA formation (Waldo et al., 1996). To further investigate the role of *Pax9* in cardiovascular development, we looked at the effect of *Pax9* loss on the pharyngeal arch transcriptome at E9.5 (**Fig. 5A-C**). Analysis of the RNA-seq data identified a total of 3863 significant differentially expressed genes (adjusted *p* value <0.1; 2073 genes up and 1790 genes down). Interestingly, *Tbx1* was significantly reduced in *Pax9^−/−^* embryos at E9.5, and this was confirmed by qPCR (**Fig. 5H**). We also confirmed that *Pax9* expression was reduced in Tbx1-null embryos (**Fig. S2**). As *Tbx1* expression is crucial for 4^th^ PAA morphogenesis (Lindsay et al., 2001; Zhang et al., 2005) this data suggested that *Pax9* may function together with *Tbx1* rather than being downstream of it. We therefore compiled a list of all the genes reported to be significantly differentially expressed in Tbx1-null embryos from published microarray and mouse genetic interaction studies (Ivins et al., 2005; Liao et al., 2008; van Bueren et al., 2010) giving a total of 1476 unique Tbx1-related genes. These were then compared to all genes significantly changed in *Pax9^−/−^* embryo pharyngeal arch tissue giving 342 genes that were common to both datasets (**Fig. 5D-F**) and suggesting that *Tbx1* and *Pax9* may share a genetic network. Among these common genes were *Chd7* and *Gbx2* (**Fig. 5I, J**) which have been shown to interact with *Tbx1* in mice (Calmont et al., 2009; Randall et al., 2009).

**Figure 5.**
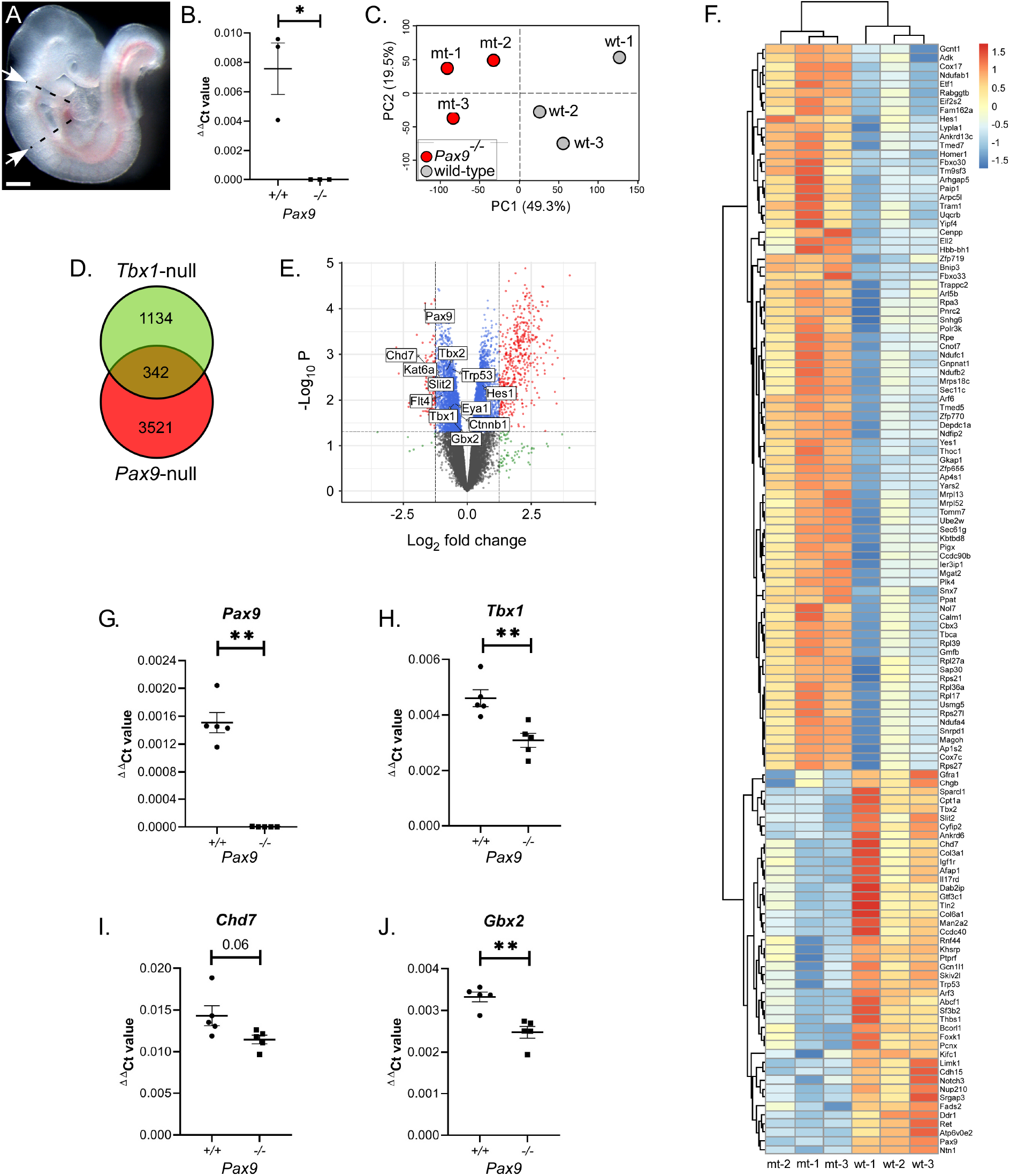
*Pax9* shares a genetic network with *Tbx1*. **A**, The pharyngeal arch region of control and *Pax9^−/−^* embryos at E9.5 (n=3 per genotype; 26-27s) was dissected free (indicated by arrows and dashed lines). Scale, 500μm. **B**, Total RNA was extracted, converted to cDNA and checked by qPCR. **C**, Principle component analysis of RNA-seq data showed a clear separation between control and mutant samples (shown on PC2) and only slight biological variation across replicates (shown on PC1). **D**, Tbx1-related genes (n=1476) were compared to those genes significantly differentially expressed in *Pax9^−/−^* embryo tissue (n=3863, adjusted *p* value <0.1) giving 342 genes common to both datasets. **E**, Volcano plot highlighting *Tbx1* interacting genes significantly differentially expressed in *Pax9^−/−^* embryos (based on adjusted *p* value ≤0.05). **F**, Heat map showing *Tbx1-Pax9* common genes that were differentially expressed (adjusted *p* value <0.05 are shown) between *Pax9^+/+^* (wt) and *Pax9^−/−^* (mt) embryos. **G-J**, qPCR analysis confirmed that *Pax9* (G), *Tbx1* **(H)**, *Chd7* **(I)** and *Gbx2* **(J)** were significantly reduced in *Pax9^−/−^* pharyngeal tissue (n=5 embryos per genotype). \*\**p*<0.01; two-tailed unpaired t-test (*Tbx1* and *Chd7*) and two-tailed Mann-Whitney (*Pax9* and *Gbx2*).

To investigate a potential genetic interaction *in vivo* between *Tbx1* and *Pax9* we confirmed that both genes were co-expressed in the pharyngeal endoderm (**Fig. S3**). We then crossed *Tbx1^+/−^* mice (Jerome and Papaioannou, 2001) with *Pax9^+/−^* mice to create *Tbx1^+/−^;Pax9^+/−^* double heterozygotes. Genotype analysis of 168 pups at weaning revealed that only two *Tbx1^+/−^;Pax9^+/−^* mice were alive, a highly significant deviation from the 42 expected of each genotype from this cross (p=1.4×10^−12^; **Table S1**). To identify the timing of lethality, neonates were observed from the day of birth with all *Tbx1^+/−^;Pax9^+/−^* mutants dying during the first 24 h. Examination of neonatal aortic arch arteries revealed that all controls had normal arch arteries (**Fig. 6A**) while *Tbx1^+/−^* neonates frequently presented with aberrant right subclavian artery (**Fig. 6B; Table 1**). All dead *Tbx1^+/−^;Pax9^+/−^* neonates had 4^th^ PAA-derived defects including interruption of the aortic arch (**Fig. 6C, D; Table 1**).

**Figure 6.**
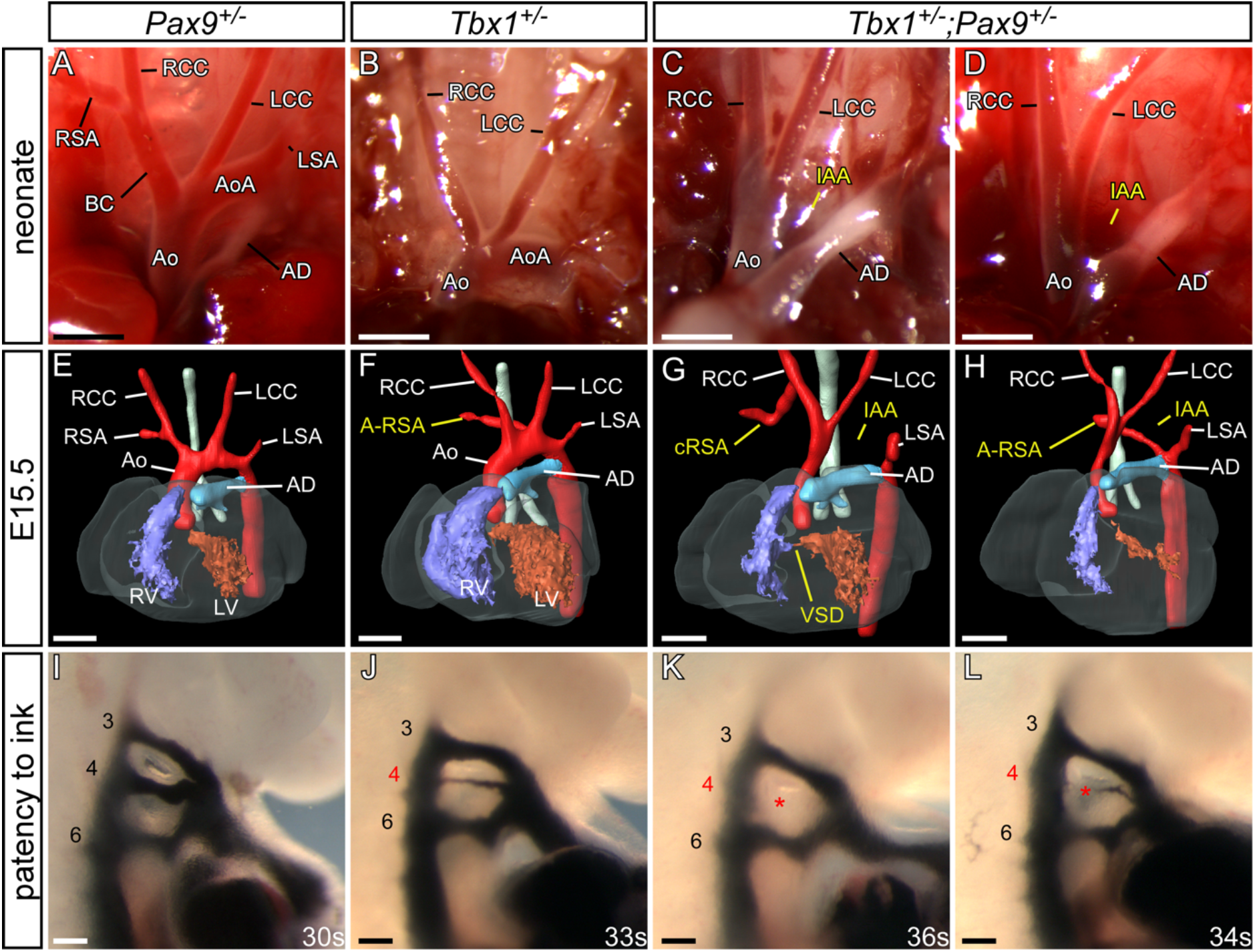
*Tbx1;Pax9* double heterozygous embryos have 4^th^ PAA defects. **A-D**, Neonates from a *Tbx1^+/−^* and *Pax9^+/−^* intercross were recovered and the aortic arches examined. **A**, Aortic arches were normal in all *Pax9^+/−^* neonates (n=10). **B**, *Tbx1^+/−^* neonates (n=6) often presented with aberrant right subclavian artery, inferred by the absence of the brachiocephalic and right subclavian arteries. **C, D**, All *Tbx1^+/−^; Pax9^+/−^* neonates (n=9) died within 24 h of birth with IAA and/or A-RSA. **E-H**, 3-D reconstructions of E15.5 embryo hearts from MRI datasets. **E**, *Pax9^+/−^* hearts were normal (n=15). **F**, *Tbx1*^+/−^ hearts (n=19) frequently displayed 4^th^ PAA defects such as A-RSA. **G, H**, All *Tbx1^+/−^; Pax9*^+/^~embryos (n=20) presented with a 4^th^ PAA-derived defect such as IAA, A-RSA and cervical right subclavian artery (cRSA). **I-L**, Intracardiac ink injections into E10.5 embryos (27-38s). **I**, PAA in control embryos was normal (n=10). **J**, In *Tbx1^+/−^* embryos (n=8) the 4^th^ PAA was often hypoplastic. **K, L**, In *Tbx1^+/−^; Pax9*^+/−^ embryos (n=9) the 4^th^ PAA was frequently bilaterally absent. Abbreviations as Figure 1. Somite counts (s) indicated. Scale: A-D, 2mm; E-H, 500μm; I-L, 100μm.

Having established postnatal cardiovascular defects we next analysed E15.5 *Tbx1^+/−^;Pax9^+/−^* embryos by MRI (**Fig. 6E-H**). All wild-type and *Pax9^+/−^* embryos were normal. Of the *Tbx1^+/−^* embryos examined, a high proportion had some form of 4^th^ PAA-derived defect, including interrupted aortic arch and aberrant right-subclavian artery (**Fig. 6F; Table 1**). All *Tbx1^+/−^;Pax9^+/−^* embryos examined, however, had some form of 4^th^ PAA-derived defect (**Fig. 6G, H; Table 1**) with the incidence of interrupted aortic arch being significantly increased compared to *Tbx1^+/−^* embryos (p<0.001). The incidence of perimembranous VSD was also significantly increased (p=0.04), but none of the embryos examined displayed outflow tract defects or bicuspid aortic valve. *Tbx1^+/−^;Pax9^+/−^* embryos at E10.5 were injected intra-cardially with ink (**Fig. 6I-L**) and the patency of the developing 4^th^ PAA was then compared between *Tbx1^+/−^* and *Tbx1^+/−^;Pax9^+/−^* embryos (Table 2). All *Tbx1^+/−^* embryos displayed a 4^th^ PAA defect, predominantly a hypoplastic vessel, consistent with previously published observations (Calmont et al., 2009; Lindsay et al., 2001; Randall et al., 2009; Ryckebusch et al., 2010). In *Tbx1^+/−^;Pax9^+/−^* embryos a significant increase in bilateral aplasia of the 4^th^ PAA (p=0.004) was seen, explaining the increase in 4^th^ PAA related cardiovascular defects observed in the *Tbx1^+/−^;Pax9^+/−^* embryos at E15.5 and in neonates (i.e. interrupted aortic arch, aberrant right subclavian artery, and cervical origins of the right subclavian artery and aorta). No defects of PAA 1, 2 and 3 were seen. This analysis therefore revealed that a concomitant heterozygosity of *Tbx1* and *Pax9* resulted in a significantly increased incidence of interrupted aortic arch caused by the failure of the 4^th^ PAA to form.

### *Tbx1* and *Pax9* interact cell autonomously in the pharyngeal endoderm for 4^th^ PAA formation

The presumed site of the *Tbx1-Pax9* interaction is the pharyngeal endoderm, and to demonstrate a cell-autonomous interaction in this tissue we crossed *Tbx1^flox^* mice (Xu et al., 2004), with a novel *Pax9Cre* transgenic mouse (**Fig. 7A-C**). This *Pax9Cre* strain has the *Cre* recombinase gene inserted into the *Pax9* coding region and effectively results in a Pax9-null allele. The *Pax9Cre* allele functions as expected by activating *R26R^lacZ^* expression specifically in the pharyngeal endoderm from E9.5 to E11.5 (**Fig. 7D-G; Fig. S4**) and *Pax9Cre;Pax9^flox^* neonates (i.e. Pax9-null) die at birth with the typical *Pax9^−/−^* phenotype (**Fig. 7H-Q**). To test the specificity of the *Pax9Cre* allele the pharyngeal arches of *Pax9Cre;eYFP* E9.5 embryos (**Fig. 8A**) were dissociated into single cells and flow-sorted into eYFP-positive and -negative populations. The eYFP-positive population was significantly enriched for *eYFP* and *Pax9* transcripts (**Fig. 8B, C**). *Tbx1^flox^* and *Pax9Cre* mice were intercrossed to create *Tbx1^+/flox^;Pax9Cre* embryos which are double heterozygous for each gene but only in the *Pax9* expression domain. A significant reduction in *Tbx1* RNA levels was observed in eYFP-positive flow-sorted pharyngeal endoderm cells from conditionally deleted E9.5 mutant embryos compared to *Pax9Cre;eYFP* controls (**Fig. 8D**). Litters from a *Tbx1^+/flox^* and *Pax9Cre* cross were collected on the day of birth, and a significant reduction in the expected number of *Tbx1^+/flox^;Pax9Cre* mutants was found (p=2.4×10^−5^; **Table S2**) and 62.5% of these mutants displayed an arch artery defect (**Fig. 8F**). We then collected embryos (E13.5-E15.5) and analysed them by MRI or μCT which revealed that 37% of mutants presented with an arch artery defect (**Fig. 8H**). We also collected embryos at E10.5 and injected them intracardially with ink to visualise the PAA. All *Tbx1^+/flox^* control embryos analysed had normal PAA, with all three vessels being patent to ink and of a similar diameter, whilst the majority of *Tbx1^+/flox^;Pax9Cre* mutant embryos presented with some form of 4^th^ PAA defect (**Fig. 8J; Table 2**).

**Figure 7.**
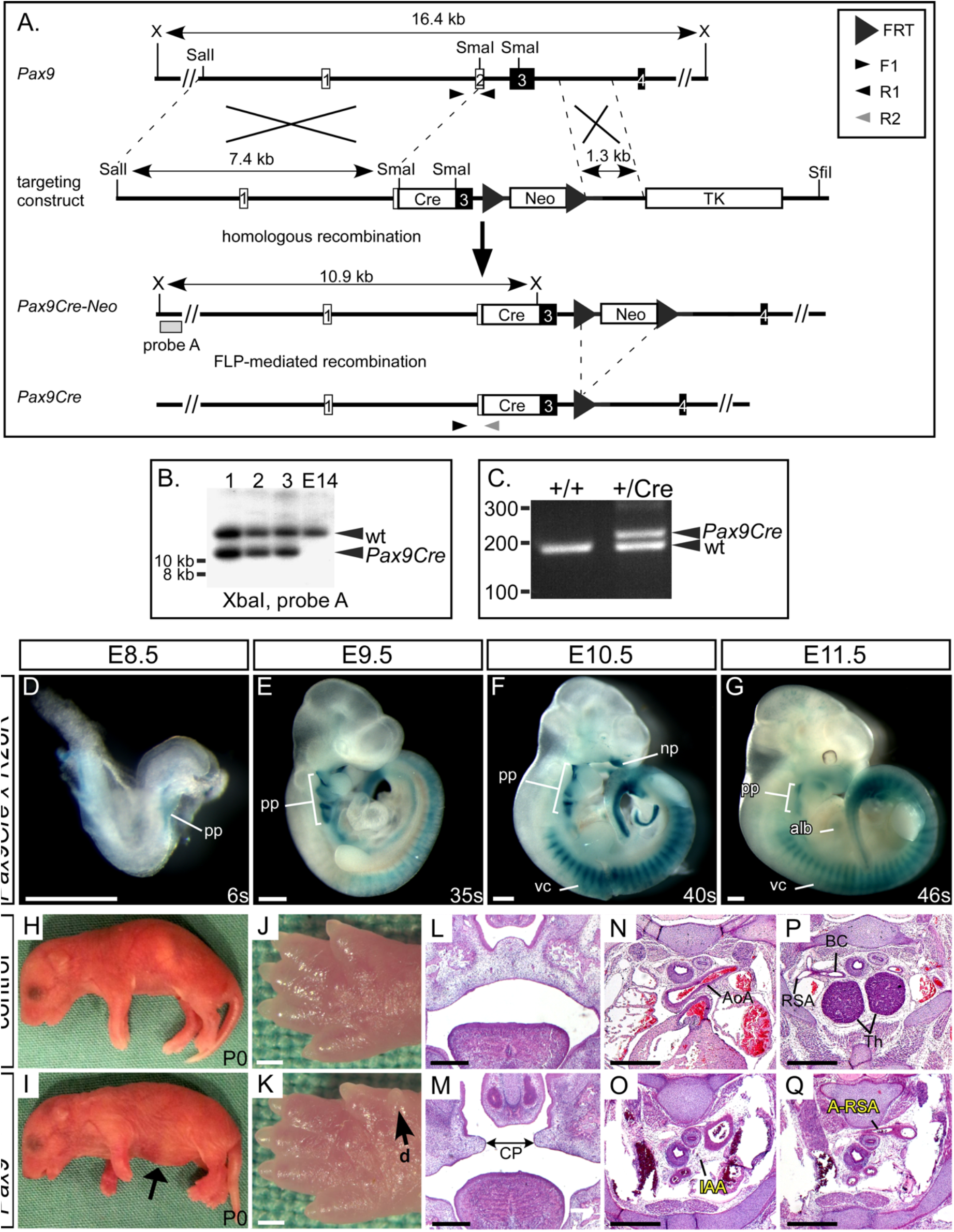
Generation of the *Pax9Cre* mouse. **A**, **A** *Pax9Cre* targeting vector was created with a *Cre* cassette inserted into the SmaI restriction sites found in exons 2 and 3 of the *Pax9* gene. For transformation the vector was linearised with SfiI. **B**, Correctly targeted ES cells were identified by Southern blotting on XbaI (X) digested genomic DNA. The derived *Pax9Cre-Neo* mice were crossed with *FLPe-* expressing mice to remove the Neomycin cassette and create *Pax9Cre* mice, which were genotyped using the indicated primers **(C). D-G**, *Pax9Cre* mice were mated with *R26R^lacZ^* reporter mice and embryos were stained with X-Gal at E8.5 (n=3), E9.5 (n=7), E10.5 (n=6) and E11.5 (n=5). Positive staining in regions known to express Pax9 was observed (alb, anterior limb bud; np, nasal process; pp, pharyngeal pouch; vc, vertebral column). H-Q, *Pax9Cre* and *Pax9^flox/flox^* mice were crossed to create *Pax9^+/flox^* (control) and *Pax9^Cre/flox^* (mutant) mice. All *Pax9^Cre/flox^* neonates died on the day of birth and presented with the typical Pax9-null phenotype (n=5), including a bloated abdomen (I, *arrow)* and pre-axial digit duplication (d) in the hind limb (**K**, *arrow*). Sections of E14.5-E15.5 embryos (n=6) showed cleft palate (CP; **M**), absent thymus and cardiovascular defects (**O, Q**). Abbreviations as Figure 1. Somite counts (s) indicated. Scale, 500μm.

**Figure 8.**
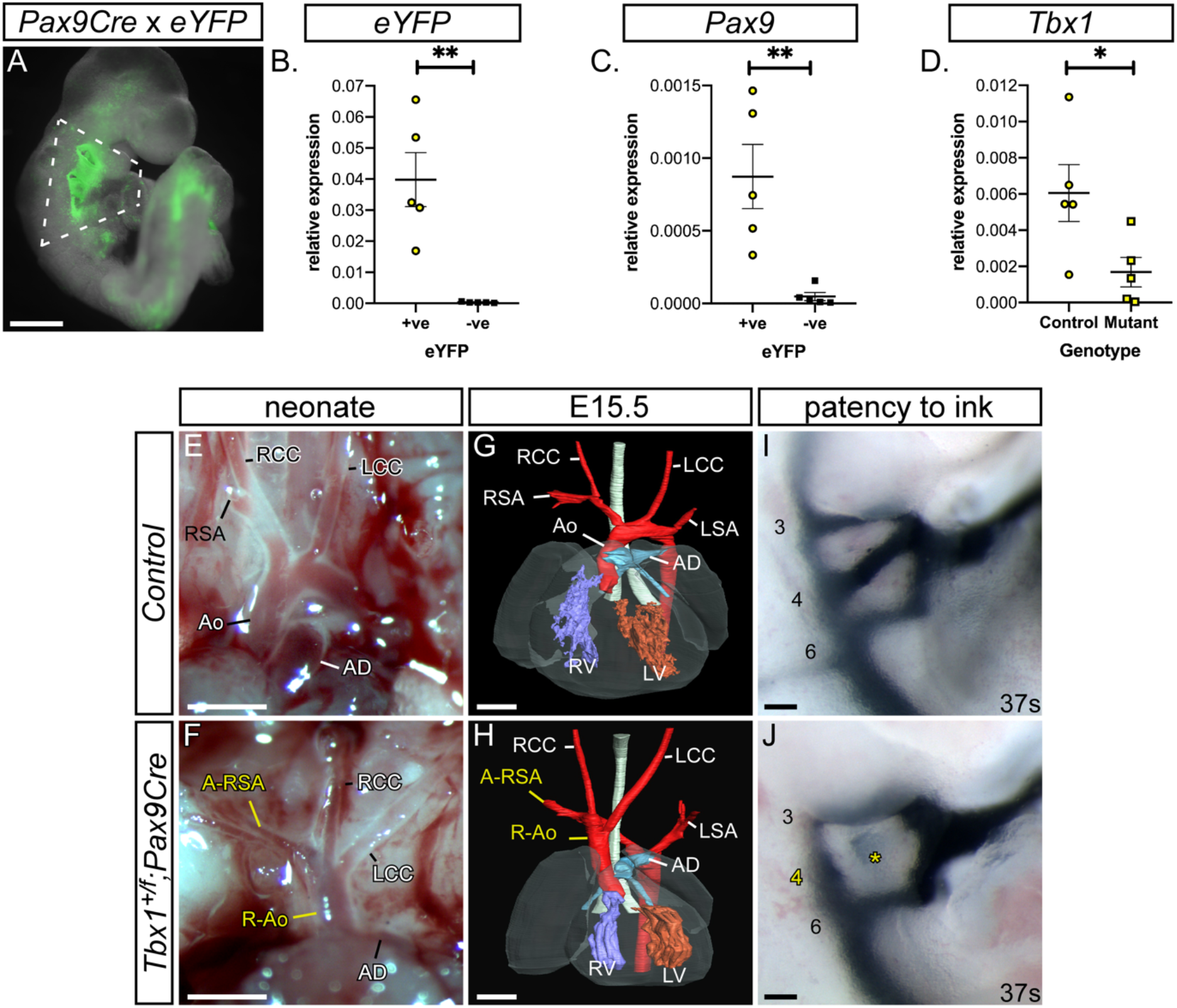
Conditional deletion of *Tbx1* from the pharyngeal endoderm. **A**, *Pax9Cre* drives expression of the eYFP reporter gene in the pharyngeal endoderm at E9.5. Scale, 500 μm. **B-D**, Flow sorting and qPCR of eYFP-positive and -negative pharyngeal arch cells from *Pax9Cre;eYFP* control and *Tbx1^+/flox^;Pax9Cre* E9.5 embryos (n=5 of each; dissected as shown in **A**). From control embryos the eYFP-positive cells *(yellow circles)* were significantly enriched for *eYFP* (**B**) and *Pax9* (**C**) compared to eYFP-negative cells *(black squares).* **D**, There was a significant reduction in *Tbx1* levels in *Tbx1^+/flox^;Pax9Cre* (mutant) eYFP-positive cells *(yellow squares)* compared to controls *(yellow circles).* *p<0.05, **p<0.01; two-tailed unpaired t-test. **E-J**, Cardiovascular defects in *Tbx1^+/flox^;Pax9Cre* mice. **E, F**, Neonates were collected shortly after birth and the aortic arch arteries examined. **E**, Control neonates (n=39 wild-type, n=43 *Pax9Cre* and n=33 *Tbx1^+/flox^)* all had normal arch arteries. **F**, *Tbx1^+/flox^;Pax9Cre* mutants (n=8) had cardiovascular defects such as a right-sided aorta (R-Ao) and A-RSA. Scale, 1mm. **G, H**, Embryos at E13.5-E15.5 were analysed by imaging (n=27 mutants). **G**, Control embryo with normal cardiovascular system. **H**, In the mutant there is a vascular ring formed by a R-Ao and an A-RSA. Scale, 500μm. **I, J**, Embryos collected at E10.5 (38-43s) were injected intra-cardially with ink. **I**, In *Tbx1^+/flox^* control embryos (n=15) PAA 3-6 were patent to ink. **J**, In *Tbx1^+/flox^;Pax9Cre* mutant embryos (n=17) the 4^th^ PAA was frequently absent. Somite counts (s) are indicated. Scale, 100μm. Abbreviations as Figure 1.

Our data, therefore, demonstrate that *Pax9* is critical for cardiovascular development and functionally interacts with *Tbx1* in the pharyngeal endoderm.

## Discussion

### Loss of *Pax9* results in complex cardiovascular defects

*Pax9* plays a key role in cardiovascular development as *Pax9^−/−^* neonates and embryos display a range of cardiovascular defects. Chromosomal deletions that include *PAX9* have been identified in patients with cardiovascular abnormalities (Hayashi et al., 2015; Kamnasaran et al., 2001; Santen et al., 2012; Schuffenhauer et al., 1999; Shapira et al., 1994) suggesting it has potential importance in human cardiovascular development. Of these, one patient with a 105 kb hemizygous deletion of 14q13 presented with interruption of the aortic arch, a hypoplastic aorta, bicuspid aortic valve, and a ventricular septal defect (Santen et al., 2012), all of which appear with high penetrance in mouse Pax9-null embryos. This small deletion only removed *PAX9, NKX2-1* and *NKX2-8*, although deletion of either *Nkx2* gene in mice does not result in any cardiovascular defects (Kimura et al., 1996; Pabst et al., 2003). It is therefore possible that hemizygous deletion of *PAX9* in humans can recapitulate, in part, the mouse knockout phenotype.

### *Tbx1* and *Pax9* interact in 4^th^ PAA development

Published microarray studies investigating the transcriptome of Tbx1-null embryos have implicated *Pax9* as a downstream target of *Tbx1*, although this was not demonstrated with any functional assays (Ivins et al., 2005; Liao et al., 2008). Analysis of our Pax9-null pharyngeal arch transcriptome data revealed that *Tbx1* levels were significantly reduced suggesting that perhaps *Pax9* and *Tbx1* may function together rather than work in a hierarchal pathway. Further analysis identified that many genes differentially expressed in Tbx1-null embryos, or shown to interact with *Tbx1* in mouse models, were also significantly differentially expressed in Pax9-null embryos, e.g. *Chd7* and *Gbx2.* Mice heterozygous for *Chd7* have abnormal 4^th^ PAA derivatives and a combined heterozygosity with *Tbx1* results in an increased incidence of 4^th^ PAA defects (Randall et al., 2009). Gbx2-null embryos display cardiovascular defects (Byrd and Meyers, 2005) and *Tbx1* and *Gbx2* have also been shown to genetically interact in PAA development (Calmont et al., 2009). Although both these *Tbx1* targets have been demonstrated to interact in the pharyngeal ectoderm, they are also expressed in the pharyngeal endoderm and could therefore interact with *Pax9* either directly or indirectly in this tissue. As our data strongly suggested that *Pax9* may interact with *Tbx1* and its related genes in the morphogenesis of the PAA we therefore looked for a genetic interaction between *Tbx1* and *Pax9* in mice. We found that all mice examined (embryos and neonates) double heterozygous for each gene (i.e. *Tbx1^+/−^;Pax9^+/−^)* displayed some defect associated with abnormal development of the 4^th^ PAA, most notably interrupted aortic arch. Although hypoplasia or absence of the 4^th^ PAA is a feature in mouse embryos heterozygous for *Tbx1* (Lindsay et al., 2001), this phenotype was significantly more prevalent in *Tbx1^+/−^;Pax9^+/−^* mice, and much higher than observed in *Tbx1^+/−^* embryos. It therefore appears that combining one Pax9-null allele with one Tbx1-null allele exacerbates the *Tbx1^+/−^* phenotype.

We used our novel *Pax9Cre* mouse to conditionally delete *Tbx1* from the pharyngeal endoderm, whilst simultaneously creating double haploinsufficiency for the two genes in this tissue. A highly significant reduction in the number of *Tbx1^+/flox^;Pax9Cre* neonates born was observed indicating that heterozygous loss of *Tbx1* and *Pax9* from the pharyngeal endoderm has a major impact on post-natal survival, with almost two-thirds of the recovered neonates examined having identifiable cardiovascular defects. A similar proportion of affected mutants was found at foetal stages. When embryos were examined at mid-embryogenesis, however, we found that almost all displayed 4^th^ PAA defects. When compared to our *Tbx1^+/−^;Pax9^+/−^* embryo data it was apparent that there was no significant difference in the incidence of bilateral 4^th^ PAA defects between *Tbx1^+/flox^;Pax9Cre* and *Tbx1^+/−^;Pax9^+/−^* embryos. This indicates that the conditional deletion of *Tbx1* from the pharyngeal endoderm, in the context of *Pax9* heterozygosity, has the same effect on 4^th^ PAA morphogenesis as seen in the global *Tbx1^+/−^;Pax9^+/−^* embryos at E10.5. Although the ink injection data from *Tbx1^+/flox^;Pax9Cre* embryos matches that seen in *Tbx1^+/−^;Pax9^+/−^* embryos, and shows that heterozygous loss of *Tbx1* from the pharyngeal endoderm severely affects PAA morphogenesis, there is a discrepancy in the cardiovascular phenotype observed at foetal and neonatal stages that does not reflect that seen at mid-embryogenesis, nor that observed in *Tbx1^+/−^;Pax9^+/−^* embryos. It is well documented, however, that the *Tbx1^+/−^* 4^th^ PAA phenotype does recover during development (see **Table S3**). It therefore appears that a conditional deletion of *Tbx1* from the pharyngeal endoderm in the context of *Pax9* haploinsufficiency replicates the *Tbx1* heterozygous arch artery phenotype in terms of a highly penetrant 4^th^ PAA defect at mid-embryogenesis that partially recovers by the foetal stage. Our data, however, does raise the question as to whether we are seeing the effect of a pharyngeal endoderm specific loss of *Tbx1* with *Pax9Cre*, or a genetic interaction occurring between *Tbx1* and *Pax9* in the pharyngeal endoderm. Conditional deletion experiments have been performed in transgenic mice to unpick the role of *Tbx1* in specific tissues and have examined the effect of *Tbx1* heterozygosity in the pharyngeal mesoderm and/or epithelium for 4^th^ PAA development (see **Table S4**). When compared, our data clearly shows that the 4^th^ PAA phenotype is much more penetrant than that seen in embryos only heterozygous for *Tbx1* in the pharyngeal epithelia. We therefore conclude that we are showing a strong genetic and cell-autonomous interaction between *Tbx1* and *Pax9* within the pharyngeal endoderm which impacts on 4^th^ PAA morphogenesis. Moreover, and in support of a phenotype extending beyond what is expected from a *Tbx1* heterozygous phenotype, our *Tbx1^flox^;Pax9Cre* mice display a vascular ring defect and a high incidence of perinatal lethality, features not previously described in *Tbx1^+/−^* mice.

### Pax9 signalling from the pharyngeal endoderm is required for 3^rd^ PAA maintenance

The mammalian PAA develop symmetrically in a cranial to caudal sequence, and then subsequently remodel to form the typical asymmetric aortic arch artery system seen in the adult (Bamforth et al., 2013; Hiruma et al., 2002). In normal PAA development the 1^st^ artery forms first, followed by the 2^nd^, but these two vessels rapidly remodel and contribute to capillary beds and the maxillary and stapedial arteries. In some *Pax9^−/−^*embryos, these vessels aberrantly persisted into E10.5 and E11.5 and may contribute to the anomalous arrangement of the aortic arch arteries when the 3^rd^ PAA collapses. In normal development the 3^rd^ PAA will form part of the common and proximal internal carotid arteries, which elongate as the embryo grows. The carotid duct involutes and the dorsal aorta anterior to this segment will form the distal internal carotid artery, with the external carotid artery derived from the proximal parts of the 1^st^ and 2^nd^ PAA (Hiruma et al., 2002). The persistence of the 1^st^ and/or 2^nd^ PAA in *Pax9^−/−^*embryos, coupled with the failure of the 3^rd^ PAA to be maintained and the carotid duct to involute, results in the external carotid arteries rising directly from the aorta, and the internal carotid arteries from the anterior dorsal aortae. This phenomenon has been described clinically (Roberts and Gerald, 1978). Although migration, apoptosis or proliferation of NCC did not appear to be impaired in Pax9-null embryos, it is well recognised that NCC differentiate into the smooth muscle cells that are important for the stabilisation of the remodelled PAA (Hutson and Kirby, 2003; Losa et al., 2017) and it is the failure of smooth muscle cells to envelop the 3^rd^ PAA in Pax9-null embryos that results in the collapse of this vessel. This has also been described in *Hoxa3-null* embryos, where smooth muscle cells fail to develop around the 3^rd^ PAA which degenerate at E11.5 (Kameda, 2009) and reduced NCC migration in genetic mouse and surgical chick models also leads to defects of the 3^rd^ PAA (Bradshaw et al., 2009; Epstein et al., 2000; Franz, 1989; Nishibatake et al., 1987). Signalling between the pharyngeal endoderm and NCC does occur (Graham et al., 2005) so it is therefore possible that signals emanating from the pharyngeal endoderm under Pax9 control are necessary to influence the differentiation of NCC into smooth muscle cells, and without these signals the 3^rd^ PAA is not supported during the remodelling phase of arch artery development and it collapses. The double outlet right ventricle phenotype observed at E15.5 in Pax9-null embryos probably arises from the marked outflow tract rotation defect due to a change in the aortic to pulmonary valve axis angle (Bostrom and Hutchins, 1988; Lomonico et al., 1986) and, possibly, aberrations in the NCC that migrate to the outflow tract may contribute to this phenotype (Bradshaw et al., 2009).

In summary, Pax9 expression in the pharyngeal endoderm has regional (anterior/posterior) roles in regulating the morphogenesis of the PAA (**Fig. 9**). In the 3^rd^ pharyngeal arch Pax9 is important for controlling NCC differentiation into the smooth muscle cells that support the remodelling vessel, whereas in the 4^th^ pharyngeal arch endoderm, the interaction of *Pax9* with *Tbx1* and its targets is crucial for the formation of the 4^th^ PAA.

**Figure 9.**
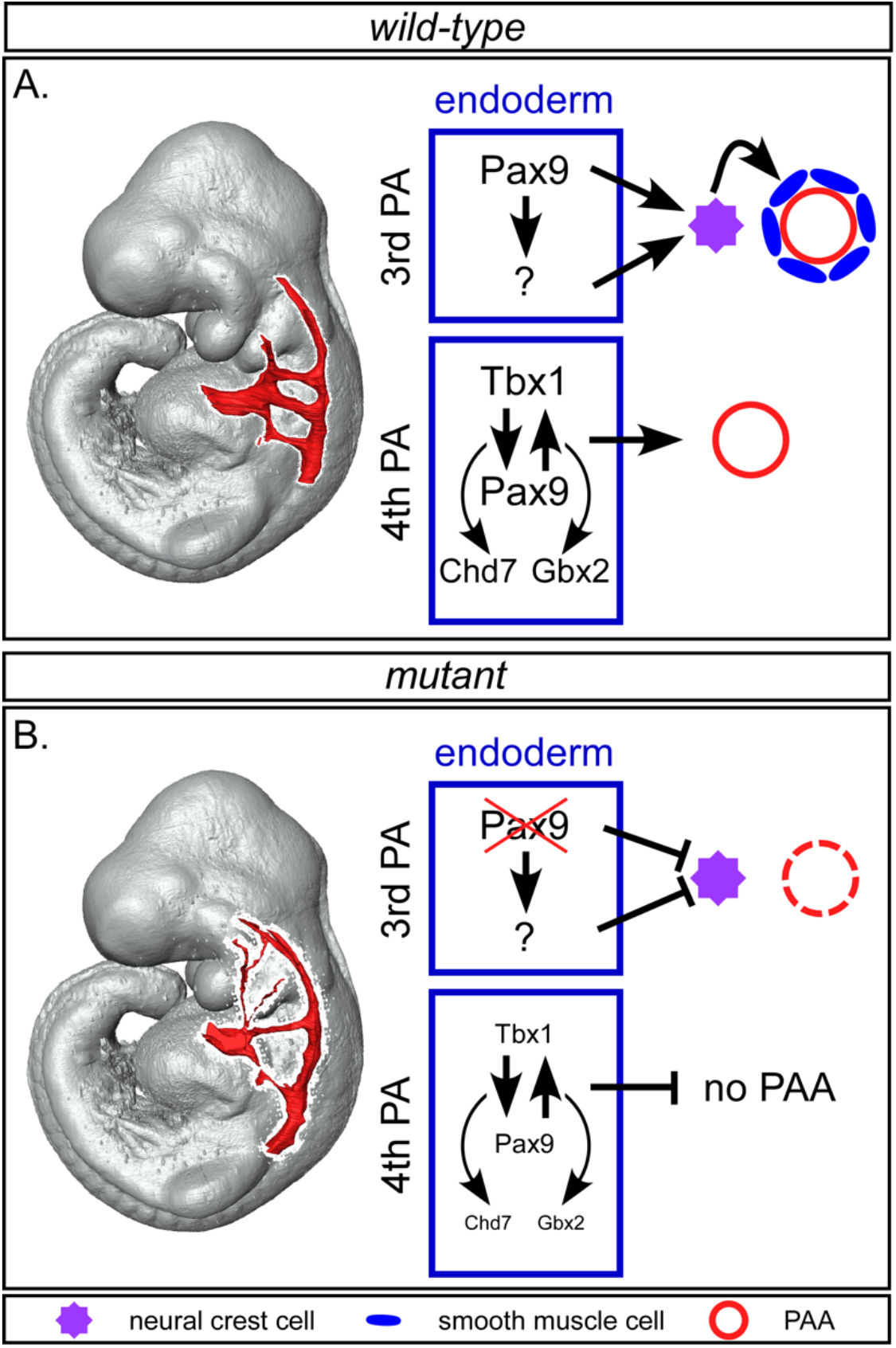
Proposed model for role of Pax9 mediated signalling during pharyngeal arch artery development. **A**, In wild-type embryos *Pax9* expression in the 3^rd^ pharyngeal arch endoderm induces neural crest cells (NCC), either directly or indirectly, to differentiate into the smooth muscle cells that invest around the endothelium of the remodelling 3^rd^ PAA. In the 4^th^ arch endoderm, *Tbx1* and *Pax9* function together to control formation of the 4^th^ PAA, potentially through collaboration with *Chd7* and *Gbx2.* **B**, In *Pax9* deficient embryos, signals from the 3^rd^ pharyngeal arch endoderm are compromised and NCC differentiation into smooth muscle cells is impaired resulting in the collapse of the 3^rd^ PAA by E11.5. In *Tbx1-Pax9* double heterozygous embryos, reduced levels of *Tbx1* and *Pax9* in the 4^th^ pharyngeal arch endoderm are insufficient to allow the 4^th^ PAA to form.

## Acknowledgements

We thank Jessica Addison, Kathleen Allinson and Divya Venkatesh for technical assistance, Sushma Grellscheid for facilitating the RNA-seq experiment, and Nicoletta Bobola for critically reading the manuscript. *Tbx1^+/−^* mice were obtained from Robert Kelly and Virginia Papaioannou. We acknowledge the Newcastle University Flow Cytometry Core Facility (FCCF) for assistance with the generation of Flow Cytometry data.

## Author contributions

CAS, WMSQ, JABL, RRK, RS, SM, RD, TB, RK: performed experiments

AIK, SC: performed RNA-seq data analysis

KS, MN, RM, RK, HP: created the *Pax9Cre* mouse

JES, TJM: generated imaging data

HMP, RK, HP: supervised the project and critiqued the manuscript

SDB: conceived and planned the experiments, supervised the project, analysed the data, created figures and wrote the manuscript

## Materials and methods

### Mice

The mice used in this study have previously been described: *Pax9^+/−^* (Peters et al., 1998), *Pax9^flox^* (Kist et al., 2007), *Tbx^+/−^*(Jerome and Papaioannou, 2001), *Tbx1^flox^* (Xu et al., 2004), *Wnt1Cre* (Danielian et al., 1998), *R26R^eYFP^* (Srinivas et al., 2001), *R26R^lacZ^* (Soriano, 1999) and *FLPe* (Dymecki, 1996). All mice were maintained on a C57Bl/6J genetic background. All studies involving animals were performed in accordance with UK Home Office Animals (Scientific Procedures) Act 1986.

### Generation of *Pax9Cre* mice

The *Pax9Cre* targeting construct was designed to replace a 1.06 kb section of the *Pax9* gene, spanning the second half of exon two, containing the start codon, and the first half of the 3^rd^ exon, containing the paired box and octapeptide motif, with a promoterless *Cre* recombinase gene preceded by a consensus Kozak sequence, in the pPNT4 targeting vector (Conrad et al., 2003), using standard molecular biology methods (**Fig. 7A**). The 1.3kb right homology arm, generated as previously described (Kist et al., 2005), was inserted between the FRT-flanked PGK-Neomycin cassette and the HSV-Thymidine Kinase cassette of pPNT4. The unwanted loxP site of pPNT4 was removed by XbaI partial digestion, followed by Klenow treatment and re-ligation. The left homology arm was generated as a 7.4 kb SalI to SmaI fragment of the *Pax9* genomic region including exon 1 and the first half of the second exon, and inserted 5’ of the *Cre* recombinase gene. The targeting vector was linearized with SfiI and mouse ES cells were targeted and selected as previously described (Kearns et al., 2013). Southern blot analysis was performed according to standard methods following XbaI digest of genomic DNA. An external southern probe was used to verify integration of the targeting cassette in the 5’ region (**Fig. 7B**) and PCR was used to validate integration of the 3’ region (not shown). The FRT-flanked PGK-Neomycin cassette was removed by crossing the derived *Pax9Cre-Neo* mice with *FLPe* mice (Dymecki, 1996) to create *Pax9Cre* mice, which were then backcrossed to strain C57Bl/6J for >6 generations. Genotyping by multiplex PCR [primers: F1-ACTCAAGCCTCTTTCAGCCC (common), R1-TTGTTCTCACTGAGCCGGCCTGT (Pax9 exon 2), and R2-GTTGCATCGACCGGTAATGC (Cre)] were used to simultaneously identify the wild-type and *Pax9Cre* alleles (**Fig. 7C**).

### Breeding

Male and female mice were mated and the detection of a vaginal plug the next morning considered to be E0.5. Pregnant females were culled on the required day and embryos collected. Embryos at E9.5-E11.5 were staged by somite counting.

### Imaging

Magnetic resonance imaging (MRI) was performed using a 9.4T MR system (Varian, US) as previously described (Bamforth et al., 2012; Schneider et al., 2004) on embryos at E15.5. High Resolution Episcopic Microscopy (HREM) and micro-computed tomography (μCT) techniques have been described in detail elsewhere (Degenhardt et al., 2010; Geyer et al., 2009). All MRI, HREM and μCT images were converted into a volume data set and segmented using Amira software (ThermoFisher Scientific) to create 2- and 3-dimensional (3-D) images. Structures were manually outlined using the label field function of Amira and surface rendered to produce the 3-D images. Intra-cardiac ink injections were performed as described (Calmont et al., 2009). Aorta measurements were taken from the region just proximal to the point at which the ascending aorta becomes the aortic arch. Haematoxylin and eosin and X-gal staining, and *Pax9* whole-mount *in situ* hybridisation, were performed using standard techniques. *Tbx1* and *Pax9* mRNA co-expression was examined by RNA *in situ* hybridisation using RNAscope^®^ Multiplex Fluorescent v2 Assay (Advanced Cell Diagnostics, Newark, CA, USA) following the manufacturer’s instructions. Probes used: mm-Pax9-C2 (454321-C2 at 1:50 dilution) and mm-Tbx1 (481911 used directly). For proliferation assays BrdU (Sigma) dissolved in PBS was administered by intraperitoneal injection at a dose of 50mg/kg to pregnant dams one hour before embryo collection.

Immunohistochemistry was performed using standard techniques (antibody details in **Table S5**) on paraformaldehyde fixed embryo sections, and fluorescently imaged on an Axioimager (Zeiss). To assess an apoptotic index within the pharyngeal arches, control and *Pax9^−/−^* embryos at E10.5 (n=3 per genotype; 30-35 somites) were examined following immunostaining with the caspase-3 antibody. To assess a proliferative index within the NCC in the pharyngeal arches, control and *Pax9^−/−^* embryos expressing *Wnt1Cre* activated eYFP at E10.5 were examined following immunostaining with anti-BrdU and anti-GFP antibodies (n=5 for each genotype; 31-39 somites). All positively stained and DAPI positive cells within the pharyngeal arches were counted from at least three slides for each embryo, and the ratio of positively stained cells over the total number of cells calculated. For endothelial cell and smooth muscle actin immunostaining n=7 control (31-36 somites) and n=6 *Pax9^−/−^* embryos (32-35 somites) were examined.

### Transcriptome analysis

Wild-type *(Pax9^+/+^*) and Pax9-null *(Pax9^−/−^)* embryos at E9.5 (n=3 of each genotype, stage-matched for 26-27 somites) were collected in ice-cold DEPC-PBS, and the pharyngeal region, from the junction between the first and second pharyngeal arch to the base of the heart, dissected free in RNAlater (Sigma). Genotypes were identified by PCR from the embryo tails and/or yolk sac material. Total RNA was extracted using the QIAGEN Plus Micro kit with genomic DNA Eliminator columns, and total RNA eluted in 30 μl RNase-free water. RNA concentration and purity were assessed by NanoDrop spectrophotometry and Agilent Bioanalyzer with all samples producing a RIN of 10. Genomic DNA contamination was excluded by PCR for intronic regions (sensitivity 5 pg/μl). To confirm that the correct genotypes were processed, cDNA was made from 100 ng total RNA and a qPCR for *Pax9* performed (**Fig. 5B**). The total RNA samples were shipped on dry-ice to EMBLM-GeneCore (Heidelberg, Germany) for RNA-seq first-strand specific library preparation using Illumina TruSeq, and 100bp paired end reads were generated from an Illumina HiSeq 2000 Sequencer. Quality control by FastQC suggested a small amount of adaptor contamination in each sample, so raw reads were trimmed of adaptor sequence with Trimmomatic (Bolger et al., 2014). Reads shorter than 36bp after trimming were discarded, and reads whose pair had been discarded were also removed. Trimmed reads were further filtered to keep only uniquely aligned and non-redundant reads to remove a vast majority of pseudogenes caused by multi-mapping. Also, to avoid bias due to imbalanced library sizes among conditions, all samples were sub-sampled towards the smallest sample (mt-663_1). The resulting reads were aligned to the Mouse Genome (Ensembl release 81, GRCm38) using the splice aware read aligner STAR 2 (star-2.5.2a-0, 20/01/2017) (Dobin et al., 2013). Finally, aligned reads were assigned to Ensembl genes using featureCounts (Liao et al., 2014) with default parameters. These count tables were imported into R and analysed using the package LIMMA (Ritchie et al., 2015) and voom function from Bioconductor (Gentleman et al., 2004).

### Data accessibility

The RNA-seq data have been deposited in NCBI’s Gene Expression Omnibus and are accessible through GEO Series accession number GSE128087 (https://www.ncbi.nlm.nih.gov/geo/query/acc.cgi?acc=GSE18087).

### Flow cytometry

The pharyngeal arch region of E9.5 *Pax9Cre;eYFP* control (n=5; 23-27 somites) and mutant *Tbx1^+/flox^;Pax9Cre;eYFP* (n=5; 24-29 somites) embryos was dissected free as described above and dissociated to single cells with Accumax (eBioscience) by incubating at 37°C for 30 minutes. The reaction was stopped by the addition of 10% fetal calf serum (FCS), the cells washed in PBS, and resuspended in 10% FCS. Cells were stained with propidium iodide. Flourescence activated cell sorting was performed on a Becton Dickinson FACS Aria II using a 100μm nozzle and a sheath pressure of 20 psi. Single cells were gated using FSC-A vs SSC-A followed by FSC-A vs FSC-H and FSC-A vs SSC-W to remove any doublets. Live single cells were gated using propidium iodide vs FSC-A, and this population finally gated on eYFP-positive and -negative cells and sorted into collection tubes.

### Quantitative real-time RT-PCR (qPCR)

RNA, from either dissociated pharyngeal arch cells or flow-sorted cells, was extracted using Trizol reagent (Invitrogen) combined with a Purelink RNA Mini Kit (Ambion) and on-column DNase1 treatment. RNA was eluted with 30 μl RNase-free water. Total RNA was converted to cDNA using a High-Capacity cDNA Reverse transcription kit (Applied Biosystems) and random hexamers. qPCR was performed using SYBR Green JumpStart Taq ReadyMix (Sigma) using the primers listed in **Table S6**. All qPCR reactions were performed in triplicate on a QuantStudio 7 Real-Time PCR System (Thermo Fisher). Data was analysed using the comparative Ct method (Schmittgen and Livak, 2008).

### Statistical analysis

Chi-squared test was used to compare genotype frequencies of litters, and Pearson’s chi-squared test for associations was used to compare defect frequencies between the different genotypes. For analysis of qPCR data, aorta measurements, and cell counts, datasets were tested for variance using the Shapiro-Wilk test (Prism 8.01 software, GraphPad). Two-tailed unpaired t-tests were performed on normally distributed data, and a two-tailed Mann-Whitney U non-parametric test was used for not normally distributed data. Groups were considered significantly different when *p*<0.05, or *p*<0.1 for RNA-seq data. No statistical methods were used to predetermine sample size which were chosen based on previous experience to obtain statistical significance and reproducibility. No data points were excluded, and all data collected from each individual experiment were used for analysis.

